# Reversing Glial Scar Back To Neural Tissue Through NeuroD1-Mediated Astrocyte-To-Neuron Conversion

**DOI:** 10.1101/261438

**Authors:** Lei Zhang, Zhuofan Lei, Ziyuan Guo, Zifei Pei, Yuchen Chen, Fengyu Zhang, Alice Cai, Yung Kin Mok, Grace Lee, Vishal Swaminnathan, Fan Wang, Yuting Bai, Gong Chen

**Affiliations:** Department of Biology, Huck Institute of Life Sciences, Pennsylvania State University, University Park, PA 16802, USA

**Keywords:** glial scar, NeuroD1, stab injury, *in vivo* conversion, glia to neuron ratio, brain repair

## Abstract

Nerve injury often causes neuronal loss and glial proliferation, disrupting the delicate balance between neurons and glial cells in the brain. Recently, we have developed an innovative technology to convert internal reactive glial cells into functional neurons inside the mouse brain. Here, we further demonstrate that such glia-to-neuron conversion can rebalance neuron-glia ratio and reverse glial scar back to neural tissue. Specifically, using a severe stab injury model in the mouse cortex, we demonstrated that ectopic expression of NeuroD1 in reactive astrocytes significantly reduced glial reactivity and transformed toxic A1 astrocytes into less harmful astrocytes before neuronal conversion. Importantly, astrocytes were not depleted after neuronal conversion but rather repopulated due to its intrinsic proliferation capability. Remarkably, converting reactive astrocytes into neurons also significantly reduced microglia-mediated neuroinflammation. Moreover, accompanying regeneration of new neurons together with repopulation of new astrocytes, blood-brain-barrier was restored and synaptic density was rescued in the injury sites. Together, these results demonstrate that glial scar can be reversed back to neural tissue through rebalancing neuron:glia ratio after glia-to-neuron conversion.

## INTRODUCTION

The central nervous system (CNS) consists of both neurons and glial cells, forming a delicate balance to maintain normal brain functions. CNS injury is often studied in the context of either neuronal loss or glial scar, but the balance between neurons and glial cells has not been adequately addressed in previous studies. One potential reason is that generating new neurons after nerve injury in the adult mammalian CNS is rather difficult despite decades of research (Cregg et al., 2014; He and Jin, 2016; Yiu and He, 2006). Another reason is that glial scar not only serves as a physical barrier but also a chemical barrier for neuroregeneration by accumulating neuroinhibitory factors such as chondroitin sulfate proteoglycans (CSPGs) and lipocalin-2 (LCN2), as well as inflammatory cytokines such as TNFα and interleukin-1β (IL-1β) (Ferreira et al., 2015; Koprivica et al., 2005; Silver and Miller, 2004). Neutralization of the neuroinhibitory factors leads to improved axonal regeneration in mouse models of CNS injury (Bradbury et al., 2002; Sivasankaran et al., 2004). The presence of glial scar correlates well with neuroinflammation and breakdown of blood-brain-barrier (BBB), resulting in a toxic microenvironment for neuron survival (Silver and Miller, 2004). Reduction of neuroinflammation in the injury sites can effectively reduce neuronal death (Fu et al., 2015; Witcher et al., 2015). However, neural protection alone is not sufficient for functional recovery if not enough new neurons being regenerated to replenish the lost neurons.

While the neuroinhibitory role of glial scar has been well established after decades of research, a recent study reported that glial scar actually helped axon regeneration based on the observation that ablation of scar-forming astrocytes inhibited axon regeneration (Anderson et al., 2016). This controversial study caused confusion in the neuroregeneration field and triggered fresh debate regarding the precise function of glial scar during neural injury and repair (Silver, 2016). Given the essential roles of astrocytes in supporting neurons, buffering excitatory toxicity, and maintaining BBB integrity (Khakh and Sofroniew, 2015), elimination of astrocytes will inevitably disrupt the CNS homeostasis. However, it is questionable whether killing reactive astrocytes is an ideal approach to assess the double-faceted roles of reactive astrocytes in terms of both neuroprotection and neuroinhibition after injury.

In this study, we employ a different strategy to reassess glial scar after neural injury through direct conversion of reactive astrocyte into neurons, rather than killing reactive astrocytes approach. We have recently demonstrated that ectopic expression of NeuroD1 in reactive astrocytes can convert them into functional neurons inside the mouse brain (Guo et al., 2014). Such glia-to-neuron conversion can also be achieved with other neural transcription factors, such as Sox2 (Heinrich et al., 2014; Niu et al., 2013; Su et al., 2014), Ngn2 (Gascon et al., 2016; Grande et al., 2013; Heinrich et al., 2014), and Ascl1 (Liu et al., 2015; Rivetti di Val Cervo et al., 2017; Torper et al., 2015; Torper et al., 2013). However, none of these studies addressed what happened to the glial scar formation after *in vivo* glia-to-neuron conversion. Will glial cells be exhausted after conversion? Will injury become worse after conversion? Will axon regeneration be inhibited after conversion?

To answer these fundamental questions regarding *in vivo* glia-to-neuron conversion, we employed a severe stab injury model in the mouse motor cortex to investigate the broad impact of cell conversion on the microenvironment of injured brains. Different from the result of killing reactive astrocytes, converting reactive astrocytes into neurons essentially reversed glial scar back to neural tissue. Importantly, astrocytes were not depleted after neuronal conversion, but rather repopulated after conversion. Unexpectedly, ectopic expression of NeuroD1 in reactive astrocytes transformed A1 type toxic astrocytes into less reactive astrocytes. Interestingly, reactive microglia were also ameliorated and neuroinflammation was reduced following NeuroD1-mediated astrocyte-to-neuron (AtN) conversion. Furthermore, blood-brain-barrier (BBB) was restored and neuronal synaptic connections were re-established after AtN conversion. Together, we demonstrate that NeuroD1-mediated cell conversion can reverse glial scar back to neural tissue by rebalancing neuron:glia ratio after injury.

## RESULTS

### High efficiency of NeuroD1-mediated astrocyte-to-neuron conversion in a severe stab injury model

The functional role of reactive glial cells after neural injury is still controversial despite extensive research over the past decades. On one hand, reactive glial cells have long been reported to secret neuroinhibitory and neuroinflammatory factors to hamper neuroregeneration (Bush et al., 1999; Cregg et al., 2014; He and Jin, 2016; Silver and Miller, 2004; Yiu and He, 2006); on the other hand, a recent study suggests that glial scar actually aids axonal regeneration (Anderson et al., 2016), raising huge confusion in the field (Silver, 2016). We have recently demonstrated that reactive glial cells can be directly converted into functional neurons inside mouse brains by a single transcription factor NeuroD1 (Guo et al., 2014). If glial scar aids axon regeneration, will converting astrocytes result in any detrimental effects and make the injury worse? To answer this important question, we established a severe stab injury model in adult mice (3-6 months old, both gender included) and investigated the impact of NeuroD1-mediated astrocyte-to-neuron (AtN) conversion on the microenvironment of the injury areas. Specifically, we used a blunt needle (outer diameter 0.95 mm) to make a severe stab injury in the mouse motor cortex, which induced a significant tissue loss together with reactive astrogliosis in the injury sites (Supplementary Fig. 1a). As expected, the number of astrocytes increased significantly in the stab-injured areas at 10 days post stab injury (dps) (Supplementary Fig. 1a, right bar graph). This is consistent with the intrinsic proliferative capability of astrocytes following neural injury, as shown by bromodeoxyuridine (BrdU) incorporation during cell division (Supplementary Fig. 1b; BrdU+ astrocytes, 38.7 ± 2.5 %, n = 4 mice, 10 dps).

To convert glial cells into neurons, we first employed retroviruses expressing NeuroD1 in dividing reactive glial cells as previously reported (Guo et al., 2014). We confirmed that ectopic expression of NeuroD1-GFP in glial cells efficiently (90.6 ± 5.2 %) converted them into NeuN+ neurons, whereas none of the GFP-infected cells were co-labeled by NeuN in the control group (Supplementary Fig. 2, n = 4 mice). Despite high conversion efficiency, the total number of newly converted neurons after retroviral infection was limited due to the limited number of glial cells that happened to be dividing during retroviral injecting. To increase the total number of newly converted neurons in the injury sites for therapeutic repair, we developed an AAV Cre-FLEX system in order to achieve more broad viral infection and cell conversion because AAV can express target genes in both dividing and non-dividing cells (Ojala et al., 2015). Specifically, Cre recombinase was expressed under the control of astrocyte promoter GFAP (GFAP::Cre) to target astrocytes specifically. The expression of Cre will act at the loxP-type recombination sites flanking an inverted sequence of NeuroD1-P2A-mCherry under the CAG promoter in a separate AAV vector (FLEX-CAG:: NeuroD1-P2A-mCherry) (Supplementary Fig. 3). Therefore, NeuroD1 expression can be targeted to reactive astrocytes where GFAP promoter is highly active, and the subsequent Cre-mediated recombination will lead to high expression of NeuroD1 driven by a strong promoter CAG. This AAV Cre-FLEX system was proved to be very efficient as shown by wide expression of NeuroD1-mCherry in the stab-injured cortical areas (Figure 1a). Importantly, at 7 days post viral injection (dpi) (injected at 4 dps), the control mCherry AAV-infected cells were mostly GFAP+ astrocytes as expected, whereas the majority of NeuroD1-mCherry infected cells had become NeuN+ neurons (Fig. 1b). Quantitatively, we found that NeuroD1-mediated astrocyte-to-neuron conversion efficiency was 89.2 ± 4.7% (7 dpi, n = 4 mice), whereas in control group 80% of mCherry AAV-infected cells were astrocytes (Supplementary Fig. 4a). The number of NeuroD1-converted neurons in the injury areas were quantified as 219.7 ± 19.3 / mm^2^ at 14 dpi. Together, by developing an AAV Cre-FLEX system to highly express NeuroD1 in reactive astrocytes, we achieved high efficiency of astrocyte-to-neuron (AtN) conversion in stab-injured mouse cortex.

**Figure 1.**
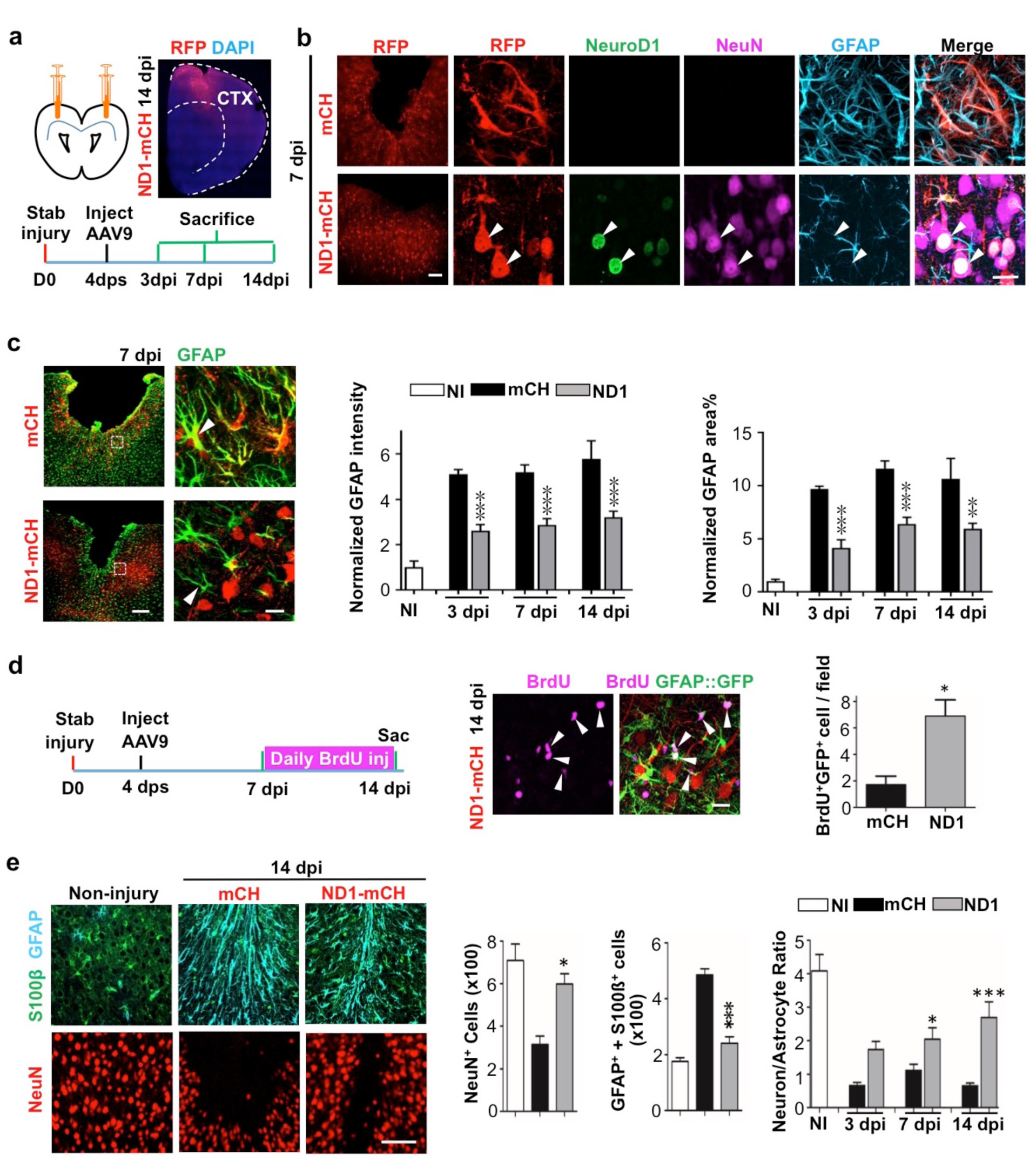
Rescue of neuron:astrocyte ratio through NeuroDI-mediated *in vivo* astrocyte-to-neuron conversion. (a)Schematic illustration showing stab injury and AAV injection in mouse motor cortex, followed by analyses at different time points. To evaluate the effect of NeuroD1-treatment, the adeno-associated virus serotype 9 (AAV9) carrying GFAP::Cre and FLEX-CAG::NeuroD1-P2A-mCherry or FLEX-CAG::mCherry-P2A-mCherry (control) were injected into the injury site at 4 days post stab injury (dps). Mice were sacrificed at 3, 7 or 14 days post viral injection (dpi) for analyses. A brain section with NeuroD1-AAV9 injection showed broad viral infection around the motor cortex area (top right). (b)NeuroD1 efficiently converted reactive astrocytes into neurons in stab-injured brains. Low-magnification images (left panels) showing the efficient viral infection in stab-injured cortex. Scale bar = 100 μm. High-magnification images (right panels) revealing the control-AAV infected cells with clear astrocytic morphology (7 dpi), while NeuroD1-AAV infected cells (arrowheads) showing neuronal morphology with high expression level of NeuroD1 (green). Co-immunostaining of NeuN and GFAP confirmed that the mCherry control-AAV infected cells (red) were GFAP+ astrocytes (top row, cyan), whereas most NeuroD1-AAV infected cells (bottom row, arrow heads) were NeuN+ neurons (magenta). Note the astrocytic morphology in the NeuroD1 group also became less hypertrophic compared to the control group. Scale bar = 20 μm. (c)Astrocytes not depleted in NeuroD1-converted areas. Control AAV-infected injury areas showed intensive GFAP signals (green) with hypertrophic morphology (top row, 7 dpi). In contrast, NeuroD1-infected injury area showed significantly reduced GFAP expression, and astrocytic morphology was less reactive but closer to healthy ones (bottom row, arrow head). Importantly, astrocytes persisted in the area with many NeuroD1-converted neurons (red). Scale bar = 200 μm (low mag), and 20 μm (high mag). Quantitative analysis revealed a significant reduction of both GFAP intensity and GFAP-covered area in NeuroD1-infected injury areas (right bar graphs). Note that the GFAP signal in NeuroD1 group was reduced to half of control group but still higher than the non-injured brains, indicating that astrocytes were not depleted after NeuroD1 conversion. n = 4-6 mice per group. ** P < 0.01, *** P < 0.001, two-way ANOVA followed by Bonferroni post-hoc test. (d)Increased proliferation of astrocytes after NeuroD1-mediated cell conversion. BrdU was applied daily in GFAP::GFP mice between 7 - 14 days post viral injection, a time window of cell conversion, to assess cell proliferation. We detected many proliferating astrocytes that were co-labeled with BrdU (magenta) and GFP (green) in the vicinity of NeuroD1-converted neurons (red). Scale bar = 20 μm. Quantitative analysis revealed a significant increase of the number of proliferating astrocytes (BrdU+/GFAP+) in NeuroD1-infected injury areas, compared to the control group. n = 3 mice. * P < 0.05, Student’s *t*-test. (e)Rescue of neuron:astrocyte ratio after NeuroD1-mediated AtN conversion. Left images illustrating neurons (NeuN, red) and astrocytes S100β, green; GFAP, cyan) in non-injured brains, mCherry control group, and NeuroD1 group. Scale bar = 100 μm. Right bar graphs, quantitative analyses illustrating the number of NeuN+ neurons, GFAP+ or S100b+ astrocytes, and the neuron:astrocyte ratio among non-injured, mCherry control, and NeuroD1 groups. Note that the neuron:astrocyte ratio was measured as 4:1 in non-injured mouse motor cortex, but significantly decreased to 0.6 after stab injury, and then reversed back to 2.6 by NeuroD1-mediated AtN conversion. n = 3-6 mice per group. * P < 0.05, *** P < 0.001, two-way ANOVA plus Sidak’s test.

### Astrocytes not depleted after conversion

Astrocytes play a vital role in supporting neuronal functions (Allen and Barres, 2009). The high efficiency of AtN conversion raises a serious concern regarding whether astrocytes might be depleted after neuronal conversion. We therefore performed GFAP staining and did observe an overall reduction of GFAP signal in the NeuroD1 group compared to the control group (Figure 1c). Unexpectedly, not only the GFAP signal was reduced, but also the astrocyte morphology showed a significant change in the NeuroD1 group, displaying much less hypertrophic processes compared to the control group (Figure 1c, arrowhead in left images), suggesting that astrocytes in the NeuroD1-converted areas became less reactive. Quantitative analysis found that the GFAP signal was significantly increased by 5-10 fold after stab injury in the control group (Figure 1c, black bar), but reduced significantly by half in NeuroD1-infected areas (Figure 1c, gray bar). The decrease of GFAP signal in NeuroD1 group is consistent with the conversion of reactive astrocytes into neurons. On the other hand, our detection of a significant level of GFAP signal in the NeuroD1-infected areas suggests that reactive astrocytes are not depleted after conversion. Since astrocytes have intrinsic capability to proliferate, we wondered whether neuronal conversion might trigger the remaining astrocytes to proliferate. To test this idea, we injected BrdU, which can be incorporated into DNA during cell division, daily from 7 dpi (viral injection at 4 dps) to 14 dpi in order to monitor cell proliferation in both control group and NeuroD1 group (Figure 1d, left schematic illustration). Interestingly, we discovered that the number of BrdU-labeled astrocytes in the NeuroD1 group more than tripled that of the control group (Figure 1d, right panels and bar graph for quantification). Importantly, many of the BrdU-labeled astrocytes were adjacent to the NeuroD1-converted neurons (Figure 1d, arrow head), suggesting that astrocytes can self-regenerate following AtN conversion. Therefore, astrocytes will not be depleted by AtN conversion but rather repopulated due to their intrinsic proliferative capability (Bardehle et al., 2013; Wanner et al., 2013).

### Neuron:astrocyte ratio rebalanced after conversion

Brain functions rely upon a delicate balance between neurons and glial cells. After neural injury, neurons die but glial cells proliferate, leading to an altered neuron:glia ratio in the injury areas. This was clearly reflected in our severe stab injury model, where the number of healthy neurons (NeuN+ cells) significantly decreased after injury but the number of astrocytes significantly increased (GFAP+/s100b+) (Figure 1e). Interestingly, after NeuroD1-mediated AtN conversion, the NeuN+ neurons significantly increased but the number of astrocytes decreased (Figure 1e), a clear reversal from the injury. Quantitative analysis discovered that the neuron:astrocyte ratio in the mouse motor cortex was ∼4:1 (4 neurons to one astrocyte) in resting condition (Figure 1e, white bar in the right bar graph). After stab injury, the neuron:astrocyte ratio dropped to <1 (Figure 1e, black bar). After NeuroD1-conversion, the neuron:astrocyte ratio reversed back to 2.6 at 14 dpi (Figure 1e, gray bar). Such significant reversal of the neuron:astrocyte ratio is critical for functional recovery in injured brains.

### A1 Reactive astrocytes transformed during conversion

A recent study suggested that reactive astrocytes after injury or disease might be characterized into A1 and A2 astrocytes with different gene expression profile (Liddelow et al., 2017). If astrocytes persisted after NeuroD1-mediated conversion, are they different from the reactive astrocytes in the control group? To answer this question, we performed RT-PCR analysis of a variety of genes related to A1 astrocytes and neuroinflammation at 3 dpi, an early stage before neuronal conversion. Compared to non-injury (NI) group, stab injury caused an upregulation of the pan-reactive astrocyte genes such as *Gfap* by 37-fold (Figure 2a), which was significantly attenuated in the NeuroD1 group (One-way ANOVA, Sidak’s test, ** P < 0.01, n = 4 pairs). Lcn2, a neuroinflammation marker associated with reactive astrocytes after injury (Zamanian et al., 2012), was increased by 700-fold after stab injury, but drastically reduced in NeuroD1 group (Figure 2a, One-way ANOVA, Sidak’s test, *** P < 0.001, n = 4 pairs). In addition, genes characteristic for A1 astrocytes such as *Gbp2* and *Serping1* were upregulated by 300-900 folds after stab injury (One-way ANOVA, Sidak’s test, *** P < 0.001, n = 4 pairs), but greatly attenuated after NeuroD1 treatment (Figure 2b). This remarkable change is unexpected, because at 3 dpi after NeuroD1 infection, astrocytes have not been fully converted into neurons yet (Supplementary Figure 5a,b). Immunostaining at 3 dpi confirmed that NeuroD1 was indeed expressed in infected astrocytes (Supplementary Figure 5b; 87.4 ± 2.5 % mCherry+ cells were NeuroD1+; 92.8 ± 2.8 % mCherry+ cells were GFAP+, n = 6 mice). Nevertheless, A1 astrocytes have already been inhibited or transformed before neuronal conversion. In addition, NeuroD1 treatment appeared to increase the astrocytic genes that support neuronal functions such as *Anax2, Thbs1*, *Gpc6, and Bdnf* (Figure 2c). As a validation of our RT-PCR result, NeuroD1 overexpression was confirmed by RT-PCR analysis (Supplementary Figure 6a). A2 astrocyte genes were not changed despite a decrease of GFAP expression after NeuroD1-infection (Supplementary Figure 6b,c).

**Figure 2.**
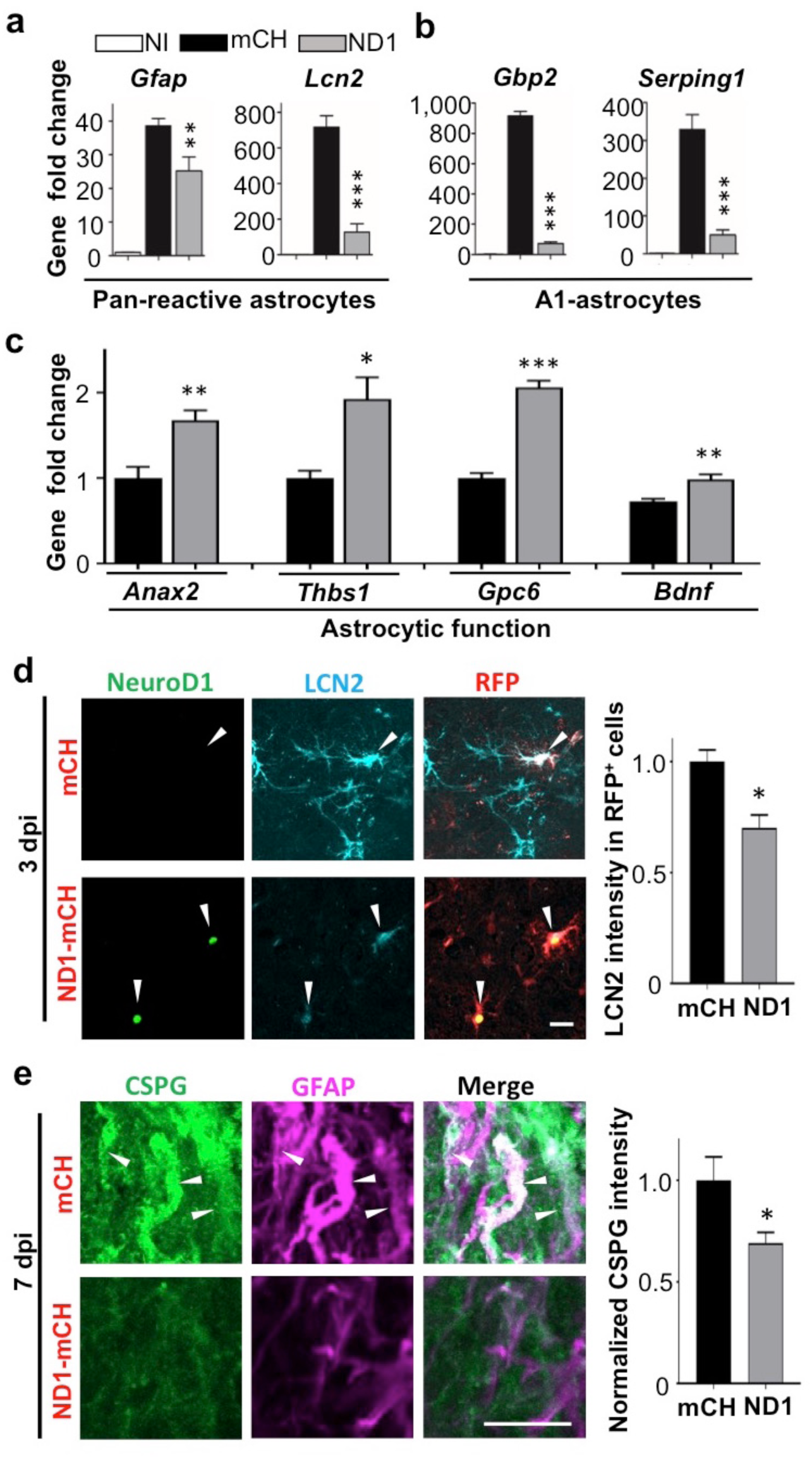
NeuroD1 transformed A1-type harmful reactive astrocytes in early time point before neuronal conversion. (a) Quantitative real-time PCR (qRT-PCR) analysis revealed a dramatic increase of reactive astrocytic genes *Gfap* and *Lcn2* after stab injury, but significantly attenuated in NeuroD1-infected areas. n = 4 mice. ** P < 0.01, *** P < 0.001, one-way ANOVA followed with Sidak’s test. (b) Toxic A1 type astrocyte-specific genes *Gbp2* and *Serping* 1 were upregulated several hundred folds in stab-injured cortices compared to non-injured cortices, but markedly reduced in NeuroD1-infected cortices. n = 4 mice. *** P < 0.001, one-way ANOVA followed with Sidak’s test. (c) More quantitative analyses with qRT-PCR revealed a significant upregulation of astrocytic functional genes in NeuroD1-infected cortices compared to the mCherry control group. * P < 0.05, ** P < 0.01, *** P < 0.001, Student’s *t*-test. (d) Representative images showing that in control-AAV infected injury sites, neural injury marker lipocalin-2 (LCN2) was highly expressed (top row, arrowhead; 3 dpi). In contrast, NeuroD1-infected cells showed much-reduced LCN2 signal (bottom row, arrowheads). Scale bar = 20 μm. Right bar graph, quantitative analysis revealed a reduced LCN2 expression in NeuroD1-infected cells compared to the control-AAV infected cells. n = 3 mice. * P < 0.05, Student’s *t*-test. (e) Representative images revealed in control-AAV infected injury sites, chondroitin sulfate proteoglycan (CSPG) was highly expressed in reactive astrocytes (upper panels, arrowheads; 7 dpi); whereas NeuroD1-infected areas showed significantly reduced CSPG signal (lower panels). Scale bar = 200 μm (low mag), 10 μm (high mag). Right bar graph showing quantitative analysis of the CSPG signal in control and NeuroD1 groups. n = 3 mice. * P < 0.05, Student’s *t*-test.

NeuroD1-positive astrocytes also showed a significant decrease in the expression level of Lcn2 compared to the reactive astrocytes in control group (Figure 2d, Student’s *t* test, * P < 0.05, n = 3 pairs), consistent with the RT-PCR analysis in Figure 2a. Furthermore, CSPG is widely associated with reactive astrocytes after neural injury and plays an important role in neuroinhibition during glial scar formation (Koprivica et al., 2005). In our severe stab injury model, we detected a high level of CSPG in the injury areas (Figure 2e, top row); but in NeuroD1-treated group, the CSPG level was significantly reduced (Figure 2e, bottom row; quantified in bar graph, Student’s *t* test, * P < 0.05, n = 5 pairs). Together, these results suggest that ectopic expression of NeuroD1 in reactive astrocytes significantly attenuated their reactive and neuroinflammatory properties. Importantly, such beneficial effects occurred as early as 3 days after NeuroD1 infection, even before astrocytes converting into neurons.

### Astrocyte-microglia interaction during neuronal conversion

It is reported that toxic A1 astrocytes are activated by cytokines such as IL-1a, TNF, and C1q secreted by reactive microglia (Liddelow et al., 2017). Conversely, reactive astrocytes also secret cytokines such as TGFb, CXCL10, CLC2, ATP, C3, ORM2 to modulate microglia (Chung and Benveniste, 1990; Norden et al., 2014). Here, we investigated what kind of impact would AtN conversion have on microglia. Compared to the resting microglia in non-injured brains (Figure 3a, top row), stab injury induced reactive microglia that were hypertrophic and amoeboid-shape (Figure 3a, middle row). As expected, both microglia and astrocytes were highly proliferative after stab injury as shown by BrdU labeling (Supplementary Figure 7a-c), but no newborn neurons detected in the adult mouse cortex after stab injury (Supplementary Figure 7d). In NeuroD1-infected areas, however, microglia morphology reversed back and was closer to the resting microglia with ramified processes (Figure 3a, bottom row). Such morphological change started as early as 3 dpi, shown in Figure 3b, where microglia contacting NeuroD1-infected astrocytes were much less reactive compared to the microglia contacting mCherry-infected astrocytes. RT-PCR analysis revealed that the cytokines TNFa and IL-1b were both significantly increased after stab injury, but both attenuated in the NeuroD1 group (Figure 3b, bar graph, One-way ANOVA followed by Turkey’s test, * P < 0.05, *** P < 0.001, n = 4 pairs). Such dramatic decrease of cytokines during AtN conversion may explain why microglia were less reactive in the NeuroD1 group. Consistently, toxic M1 microglia that were immunopositive for iNOS showed a significant reduction in the NeuroD1 group compared to the control group (Figure 3c, Student’s *t* test, *** P < 0.001, n = 4 pairs). Note that the reduction of iNOS-labeled M1 microglia coincided with the reduction of toxic A1 astrocytes, as detected at 3 dpi after NeuroD1 infection, suggesting an intimate interaction between astrocytes and microglia (Volterra and Meldolesi, 2005). At 7 dpi, compared to the control group, the reduction of iNOS in the NeuroD1 group was even more significant, accompanied with a reduction of Iba1 signal as well (Figure 3d, two-way ANOVA, ** P < 0.01, *** P < 0.001, n = 6 pairs at 3 dpi and 7dpi, n = 5 pairs at 14 dpi). In addition, stab injury also induced a remarkable increase in the expression level of CD68, a marker for macrophages and monocytes as well as some reactive microglia (Figure 3e). However, in NeuroD1-infected areas, the expression level of CD68 was significantly attenuated (Figure 3e, bar graph, two-way ANOVA, ** P < 0.01, *** P ≤ 0.001, n = 6 pairs). Together, these results suggest that accompanying astrocyte-to-neuron conversion, toxic M1 microglia are reduced and neuroinflammation is alleviated.

**Figure 3.**
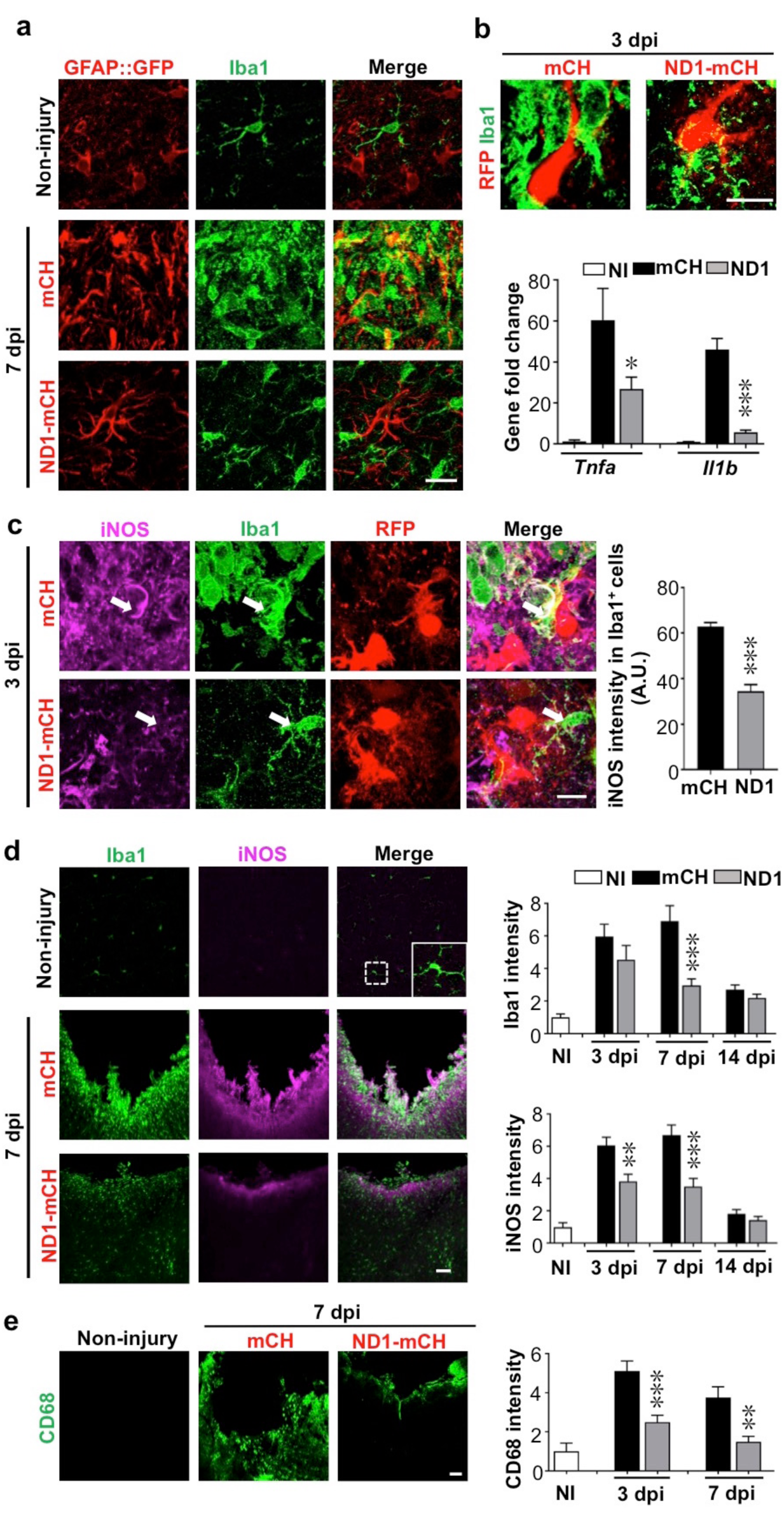
NeuroD1-treatment attenuated microglial inflammatory responses. (a)Microglia (Iba1, green) in non-injured brains displaying ramified branches (top row), but showing hypertrophic amoeboid shape in stab-injured areas (middle row, 7 dpi). In NeuroD1-infected injury areas, however, microglia returned to ramified morphology again (bottom row). Importantly, such morphological changes of microglia coincided with the morphological changes of astrocytes (GFAP::GFP labeling in left column). Scale bar = 20 μm. (b)Representative images showing a close look of the microglia morphology (Iba1, green) contacting mCherry-infected astrocytes (left panel, red) or NeuroD1-mCherry infected astrocytes (right panel, red) at 3 dpi. Note that microglia showed clear morphological difference when contacting the NeuroD1-infected astrocytes as early as 3 dpi, before neuronal conversion. Scale bar = 20 μm. Lower bar graph illustrating the gene expression level of inflammatory factors *Tnfa and Il1b* significantly increased in stab-injured cortices compared to non-injured cortical tissue, but such increase was greatly reduced in NeuroD1-infected injury areas (3 dpi). n = 4 mice. * P < 0.05, *** P < 0.001, one-way ANOVA followed with Sidak’s test. (c)Representative images illustrating many inflammatory M1 microglia labeled by nitric oxide synthase (iNOS) with amoeboid morphology in the control-AAV infected injury areas (upper panels, 3 dpi). In contrast, microglia in close contact with the NeuroD1-infected astrocytes showed much lower iNOS expression with ramified morphology (lower panels, arrow; 3 dpi). Scale bar = 10 μm. Quantitative analysis showing a significant reduction of iNOS signal of Iba1+ cells in close contact with NeuroD1-infected cells. n = 3-4 mice per group. *** P < 0.001, Student’s t-test. (d)Representative images illustrating lack of iNOS signal in non-injured brains (upper panels), but high Iba1 and iNOS signal in stab-injured cortical tissue (middle panels, 7 dpi). However, in NeuroD1-infected cortices, both Iba1 and iNOS signals reduced significantly (bottom panels, 7 dpi). Scale bar = 50 μm. Right bar graphs, quantitative analyses showing the immunofluorescent signal of Iba1 and iNOS in non-injured (white bar), mCherry control (black bar), or NeuroD1 AAV-infected cortices (gray bar) at 3, 7 and 14 dpi. ** P < 0.01, *** P < 0.001. Two-way ANOVA followed by Bonferroni post-hoc tests. n = 5-6 mice per group. (e)Representative images showing the immunoreactivity of CD68, a macrophage marker, significantly reduced in NeuroD1-infected cortical tissues. Scale bar = 50 μm. Right bar graph showing quantitative analysis result, which revealed a significant reduction of CD68 fluorescent signal in NeuroD1-infected cortices (gray bar) at 3 and 7 dpi. n = 5-6 mice per group. ** P < 0.01, *** P < 0.001, two-way ANOVA plus Bonferroni post-hoc test.

### Astrocyte-blood vessel interaction during *in vivo* cell conversion

One important function of astrocytes in the brain is to interact with blood vessels and contribute to blood-brain-barrier (BBB) in order to prevent bacterial and viral infection and reduce chemical toxicity (Obermeier et al., 2013). Breakdown of BBB will result in the leakage of a variety of biological and chemical agents into the parenchymal tissue, contributing to the secondary neural injury (Bush et al., 1999). In healthy brain, BBB is tightened by astrocytic endfeet wrapping around the blood vessels (Figure 4a). Comparing to the evenly distributed blood vessels (labeled by endothelial marker Ly6C) in non-injured brains (Figure 4b, left panels), stab injury caused blood vessels swollen (Figure 4b, middle panels). In NeuroD1-treated group, however, blood vessels exhibited less hypertrophic morphology and closer to the ones in healthy brains (Figure 4b, right panels). Accompanying altered blood vessel morphology after stab injury, we also found a disruption of BBB, as evident by the mislocalization of AQP4 signal. AQP4 is a water channel protein, normally concentrating at the endfeet of astrocytes at resting state wrapping around blood vessels (see Figure 4a). After stab injury, AQP4 signal dissociated from blood vessels and instead distributed throughout the injury areas (Figure 4c, top row). Interestingly, in NeuroD1-treated areas, AQP4 signal showed reassociation with blood vessels, returning back to normal state (Figure 4c, bottom row). Note that the astrocytic morphology also looked much less reactive in NeuroD1-treated group (Figure 4c, bottom row). To further evaluate BBB integrity, we perfused the mice with biotin, a molecule that can easily leak out after BBB breakdown. After stab injury, we observed a significant leakage of biotin in the injured areas in control group expressing mCherry alone (Figure 4d, top row). However, in NeuroD1 group, biotin was mainly detected inside the blood vessels and the leakage was significantly reduced in the parenchyma tissue (Figure 4d, bottom row), indicating restoration of BBB integrity. Together, these results suggest that after NeuroD1-mediated cell conversion, astrocytes interact with blood vessels again to restore the broken BBB caused by neural injury.

**Figure 4.**
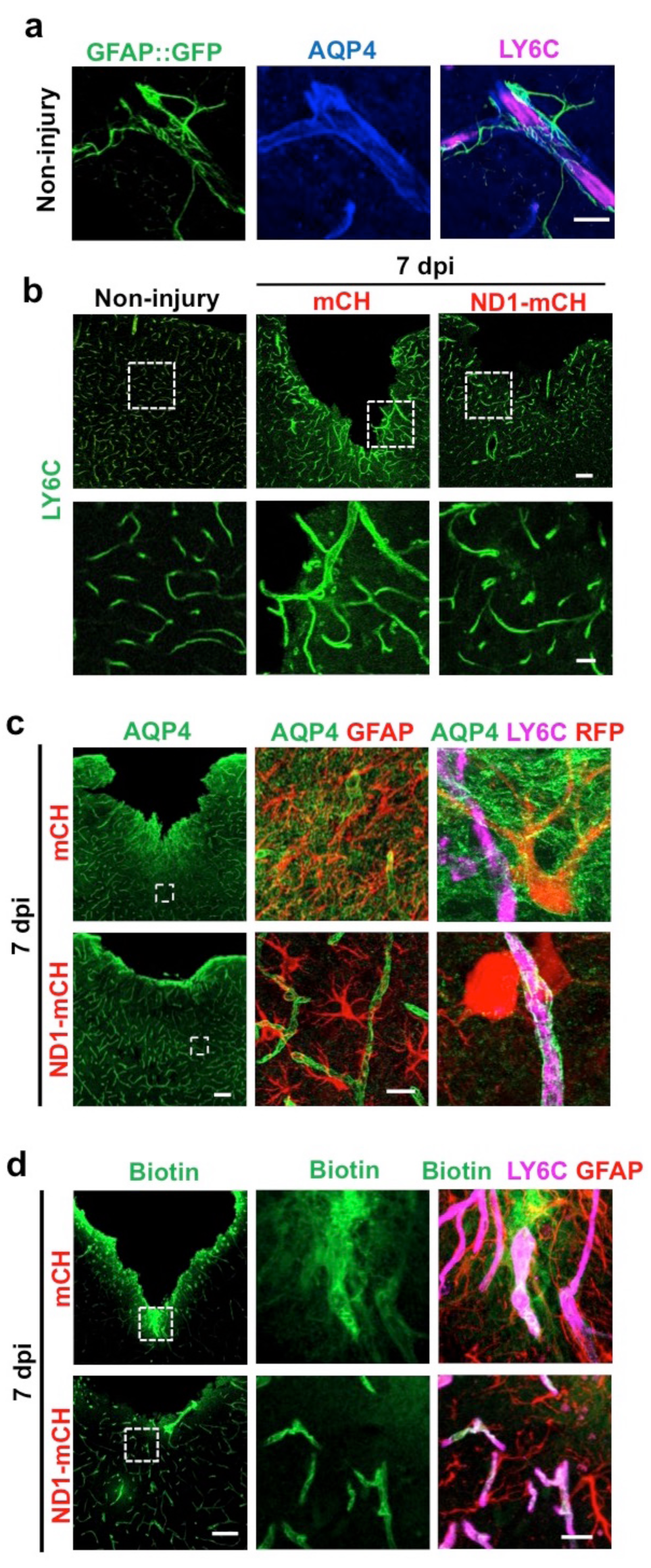
Repair of blood vessels and blood-brain-barrier after stab injury through NeuroDI-mediated *in vivo* cell conversion. (a) Representative images showing the astrocyte-vascular unit in non-injured mouse cortex. Astrocytes (green, labeled by GFAP::GFP) send their endfeet wrapping around blood vessels (magenta, labeled by LY6C, a vascular endothelial cell marker). Water channel protein aquaporin 4 (AQP4, blue) was highly concentrating at the astrocytic endfeet in resting state, which wrapped around the blood vessels. Scale bar = 20 μm. (b) Representative images in lower magnification (top row) and higher magnification (bottom row) showing the blood vessel morphology disrupted by stab injury. In NeuroD1-infected areas, the hypertrophic blood vessel morphology was partially reversed, closer to the ones in non-injured brains. Scale bar = 100 μm (low mag), 20 μm (high mag). (c) Top row illustrating mislocalization of AQP4 signal (green) and detachment of astrocytic endfeet from blood vessels (magenta, LY6C) after stab injury. Note the AQP4 signal was spreading throughout the parenchyma tissue without concentrating around the blood vessels. Bottom row illustrating in NeuroD1-infected areas, AQP4 signal (green) was re-associated with blood vessels (magenta, LY6C). Scale bars = 100 μm (low mag), 20 μm (high mag). (d) Top row illustrating a significant leakage of biotin into the parenchyma tissue after stab injury, suggesting a disruption of blood-brain-barrier (BBB) integrity (7 dpi). Note that biotin signal was found not only inside the blood vessels but also outside the blood vessels. Bottom row illustrating that in in NeuroD1-infected areas, biotin was mostly confined inside blood vessels, suggesting a restoration of BBB integrity. Scale bars = 100 μm (low mag), 20 μm (high mag).

### Functional recovery after NeuroD1-mediated neuronal conversion

With a substantial change in the glial environment after NeuroD1-mediated AtN conversion, we further investigated neuronal properties such as dendritic morphology, synaptic density, and electrophysiological function in the injury areas. Stab injury resulted in severe dendritic damage as expected, shown by dendritic marker SMI32 (Figure 5a, left images, green signal). In NeuroD1-infected areas, however, we detected a significant increase in dendritic signal SMI32 (Figure 5a, bottom image). Quantitatively, neuronal dendrites labeled by SMI32 were severely injured after stab lesion, decreasing to 25% of the non-injured level, but NeuroD1 treatment rescued the dendritic signal to over 50% of the non-injured level at 14 dpi (Figure 5a; bar graph, two-way ANOVA, *** P < 0.001, n = 4 pairs). Consistent with the dendritic damage, stab injury also caused a severe reduction of synapses in the injured areas as shown by glutamatergic synapse marker vGluT1 and GABAergic synapse marker GAD67 (Figure 5b, left images). After NeuroD1 treatment (30 dpi), both glutamatergic and GABAergic synaptic density in the injured areas showed a significant increase compared to the control group (Figure 5b, bar graphs).

**Figure 5.**
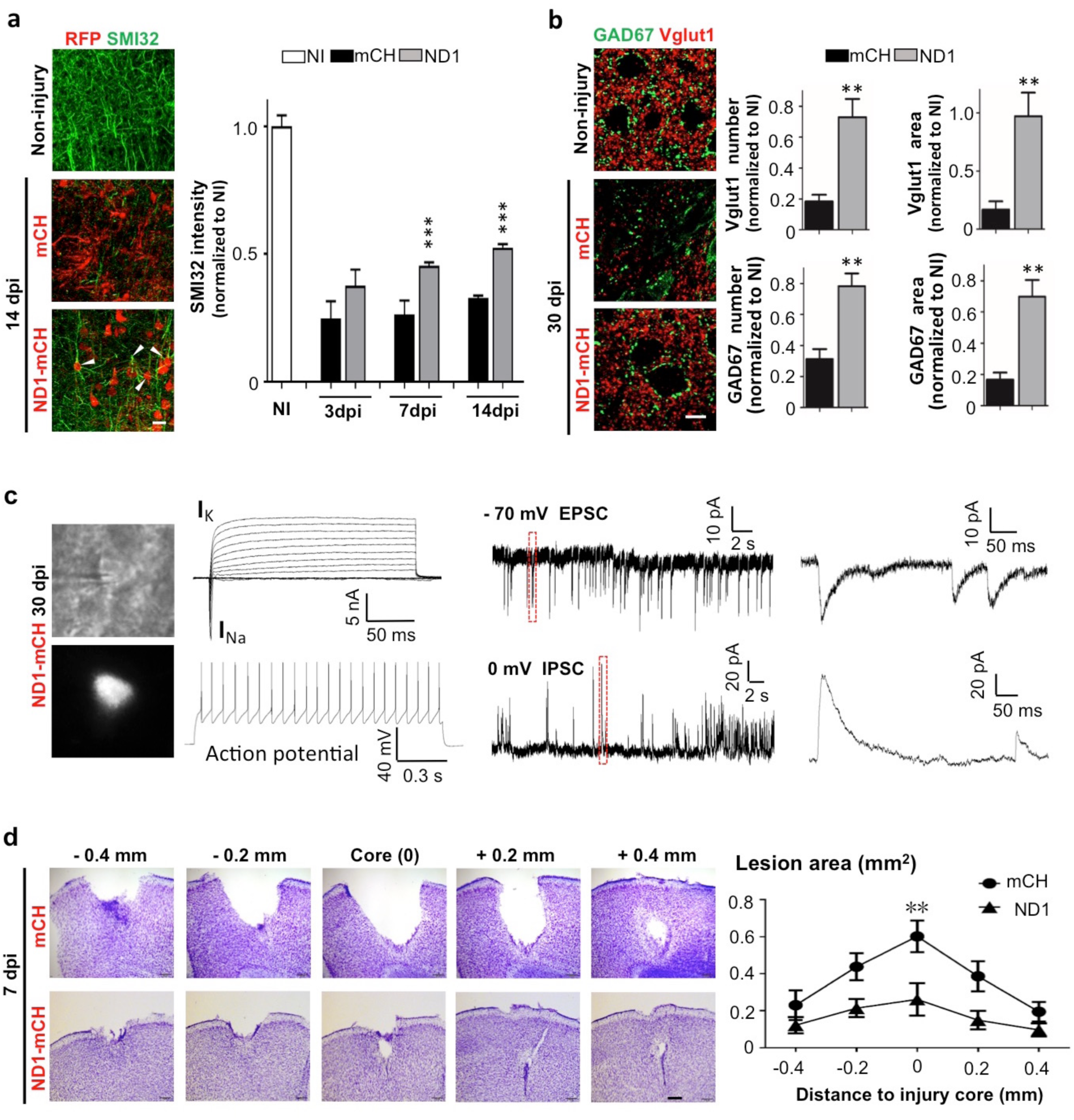
Functional rescue by NeuroD1-mediated astrocyte-to-neuron conversion. (a) Left images showing dendritic marker SMI32 (green) drastically reduced after stab injury (middle panel), but significantly rescued in NeuroD1-infected injury regions (bottom panel). Note that the NeuroD1-converted new neurons (red, bottom panel) were co-labeled by SMI32, indicating the newly generated neurons contributing to neural repair. Right bar graph illustrating SMI32 signal partially recovered at 7 and 14 dpi. Scale bar = 20 μm. n = 3-4 mice per group. *** P < 0.001, two-way ANOVA plus Bonferroni post-hoc test. (b)Rescue of synaptic loss after stab injury by NeuroD1-mediated cell conversion. Left images showing both glutamatergic synapses (red, Vglut1) and GABAergic synapses (green, GAD67) significantly reduced after stab injury (middle panel, 30 dpi). However, after NeuroD1-treatment, both glutamatergic and GABAergic synapses showed significant recovery. Scale bar = 20 μm. Right bar graphs showing quantitative analysis results. Note that both the number of synaptic puncta and the covered area were rescued in NeuroD1-treated injury areas. n = 4-5 mice per group. *** P < 0.001, two-way ANOVA plus post-hoc test. (c)Electrophysiological analysis demonstrating that the NeuroD1-converted neurons were functional. Left images showing patch-clamp recordings performed on NeuroD1-mCherry-infected cells in cortical slices (30 dpi). Right traces showing typical recording of large Na+ and K+ currents, repetitive action potentials, and excitatory (EPSC) and inhibitory (IPSC) postsynaptic currents. n = 15 cells from 3 mice. (d)Rescue of tissue loss by NeuroD1-mediated AtN conversion. Top row illustrating cortical tissue damage induced by stab injury with Nissl staining in a series of brain sections across the injury core (7 dpi). Bottom row illustrating much less tissue loss in the NeuroD1 group. Scale bar = 200 μm. Right line graph showing the quantitative analysis result. Across the entire injury areas, cortical tissue loss was significantly rescued throughout the NeuroD1-infected regions. The tissue areas absent of crystal violet signals were quantified. n = 5 mice. ** P < 0.01, two-way ANOVA followed with Bonferroni post-hoc test.

We next investigated whether the NeuroD1-converted neurons are physiologically functional. Cortical brain slices were prepared and patch-clamp recordings were performed on the NeuroD1-mCherry converted neurons (Figure 5c, left images). After one month of conversion, the NeuroD1-mCherry positive neurons showed large sodium and potassium currents (>5 nA) and repetitive action potentials (Figure 5c, left traces). More importantly, we recorded both glutamatergic synaptic events (frequency: 0.96 ± 0.5 Hz; amplitude: 24.4 ± 6.3 pA; holding potential = -70 mV) and GABAergic synaptic events (frequency: 0.74 ± 0.16 Hz; amplitude: 55.9 ± 7.7 pA; holding potential = 0 mV) (Figure 5c, right traces), consistent with the recovery of vGluT1 and GAD67 immunopuncta shown in Figure 5b.

Finally, we repeatedly observed a significant tissue loss after severe stab injury during a variety of immunostaining analysis, but a remarkable tissue repair in the NeuroD1-treatment group (Supplementary Figure 4b). To quantitatively assess the level of tissue loss, we performed serial cortical sections with Nissl staining around the injury core in both mCherry control group and NeuroD1 treatment group. Nissl staining revealed a large tissue loss across the serial brain sections in the mCherry control group after stab injury (Figure 5d, top row). In contrast, the NeuroD1-treatment group showed much less tissue loss across all brain sections (Figure 5d, bottom row). Quantitatively, the tissue loss around the injury areas in the NeuroD1 group was significantly reduced by 60% compared to the control group (Figure 5d, right line graph, Two-way ANOVA, ** P < 0.01, n = 5 pairs). Together, these results suggest that converting reactive astrocytes into neurons significantly reduced tissue loss and successfully repaired the damaged brain.

## DISCUSSION

In this study, we demonstrate that NeuroD1-mediated astrocyte-to-neuron conversion essentially reverses glial scar back to neural tissue through restoring the neuron:astrocyte ratio in the injury sites. This is achieved not only by regeneration of new neurons, but also self-repopulation of new astrocytes after conversion. Even before converting into neurons, toxic A1 type reactive astrocytes can be transformed into less reactive astrocytes and hence releasing less toxic cytokines after expressing NeuroD1. Meanwhile, toxic M1 microglia and neuroinflammation in the injury sites are attenuated, and blood vessels and BBB are restored after NeuroD1-mediated AtN conversion. Together, AtN conversion results in more functional neurons, less reactive glial cells, and rebalanced neuron:astrocyte ratio. Therefore, it is possible to reverse glial scar back to neural tissue through *in vivo* convertion of reactive astrocytes into functional neurons.

### Reversing glial scar back to neural tissue through rebalancing neuron:glia ratio

Glial cells, in particular astrocytes, closely interact with neurons to provide structural, metabolic, and functional support. A delicate balance between neurons and glia ensures normal brain function. Injury to brain tissue leads to irreversible neuronal loss accompanied by glial proliferation. As a result, the delicate balance between neurons and glia is altered and brain functions impaired. In the mouse motor cortex where our studies are performed, the neuron:astrocyte ratio was quantified to be 4:1 (4 neurons/astrocyte), which is in accordance with previous findings (Nedergaard et al., 2003). Considering its long axon and dendrites, each individual neuron is actually associated with many astrocytes. Although the functional role of astrocytes in neural injury has been extensively examined in previous studies, the balance between neurons and glia during brain repair has not been well understood. This may be partly due to the fact that previous technologies are not possible to restore the neuron:glia ratio after injury. Transplantation of exogenous neuroprogenitor cells can generate new neurons, but difficult to reduce reactive glial cells (Lu et al., 2012; Peron et al., 2017; Toft et al., 2007). Conversely, ablation of astrocytes can reduce the number of glial cells but cannot generate new neurons (Anderson et al., 2016). Other approaches attempting to attenuate glial scar, such as using chondroitinase ABC to reduce CSPG level (Sekiya et al., 2015), also cannot generate new neurons. We predict that brain functional recovery will be difficult to achieve if neuron:glia ratio after injury is not restored to a level close to the resting state.

Comparing to previous technologies, our *in vivo* AtN conversion approach offers a unique strategy that can both reduce reactive astrocytes without killing them, and generate sufficient number of new neurons without external cell transplantation. By directly converting reactive astrocytes into neurons, we are able to quickly rebalance neuron to astrocyte ratio from 0.6 after brain injury to 2.5 in 2 weeks after neuronal conversion, which is much closer to 4 neurons/astrocyte in the resting state. Accompanying such remarkable increase of neuron:astrocyte ratio, we demonstrate that neuronal dendrites are significantly restored together with a significant increase of synaptic density. Therefore, we conclude that *in vivo* cell conversion can rebalance neuron:glia ratio and reverse glial scar back to functional neural tissue.

### Astrocytes not depleted but rather repopulated after neuronal conversion

As the most abundant cell type in the mammalian CNS, astrocytes influence brain functions by providing trophic support to neurons, regulating BBB, modulating immune cells, and controlling synapse formation. It is well established that reactive astrocytes undergo dramatic morphological change and express high level of GFAP and cytokines such as TNFa and Lcn2 in the CNS after injury or disease (Zamanian et al., 2012). However, the precise function of reactive astrocytes after injury is still under debate due to their double-faceted roles. On one hand, the inhibitory effect of reactive astrocytes on axon regeneration has been widely reported (Cregg et al., 2014; Silver and Miller, 2004; Yiu and He, 2006). Reactive astrocytes also produce pro-inflammatory cytokines that exacerbate CNS injury (Brambilla et al., 2005; Pineau et al., 2010; Sofroniew, 2015). On the other hand, a recent study reported that ablation of scar-forming astrocytes after spinal cord injury impaired axon regeneration, leading to a conclusion that glial scar actually aids axon regeneration (Anderson et al., 2016). In line with this finding, the same group earlier reported that the depletion of reactive astrocytes led to spreading of a stab wound in the forebrain, indicated by prolonged immune response, failure of BBB repair, and extensive neural degeneration (Bush et al., 1999). It is important to point out that these results are based on the ablation of scar-forming astrocytes, which causes massive astrocytic death in the injury areas and will inevitably trigger massive neuroinflammation from reactive microglia. This is indeed what has been observed when astrocytes are killed (Anderson et al., 2016; Bush et al., 1999). Over activation of microglia and macrophages can induce profound neurotoxicity and axonal retraction through both physical interaction and inflammatory responses (Horn et al., 2008; Hu et al., 2015; Silver, 2016). In addition, astrocytes provide a variety of support to normal neuronal functions, ranging from neurotrophic nourishing to neurotoxin removal as well as regulating synaptic structure and functions (Allen and Barres, 2009). Therefore, it is not surprising that killing astrocytes will further deteriorate the microenvironment of injury areas if neurons are deprived of astrocytic support. In contrast to the killing astrocyte approach, we demonstrate here that our *in vivo* neuronal conversion approach did not deplete astrocytes in the injured areas. Instead, after AtN conversion, the remaining astrocytes showed much higher proliferation rate to repopulate themselves. Therefore, different from astrocyte-killing approach, our *in situ* AtN conversion approach not only regenerates new neurons but also regenerate new astrocytes, reaching a new balance between neurons and astrocytes in the injury areas.

Why killing astrocytes results in worsening of neural injury, whereas converting astrocytes into neurons repairs damaged neural tissue? The fundamental difference is the way astrocytes are treated. When transgenic mice were used to kill proliferating reactive astrocytes or impair STAT3 signaling pathway, not many functional astrocytes were left in the injury areas (Anderson et al., 2016). In contrast, when we converted reactive astrocytes into neurons, we observed many astrocytes with less reactive morphology in the injury areas. While this was initially surprising to us, it later became more apparent after we discovered that the entire microenvironment was ameliorated after AtN conversion. One of the new insights gained from this study is that in addition to neural injury that can stimulate astrocytic proliferation, we now demonstrate that AtN conversion in the adult brain can also trigger astrocytic proliferation. With repopulation of astrocytes in the injury areas, our AtN conversion approach avoids the detrimental effects of depleting astrocytes reported before (Anderson et al., 2016). In fact, we believe that the newly generated astrocytes may interact with the newly converted neurons to facilitate neural repair.

Besides astrocytic proliferation after conversion, another unexpected finding is the transformation of A1-like toxic reactive astrocytes into non-toxic astrocytes after ectopic expression of NeuroD1 before neuronal conversion. Recent report identified A1 toxic reactive astrocytes after CNS injury (Liddelow et al., 2017; Zamanian et al., 2012). Specifically, A1 astrocytes lose normal astrocytic functions but exhibit detrimental effects on CNS repair such as secreting neurotoxic cytokines, disrupting synapses, and killing neurons and oligodendrocytes. Our data revealed that the reactive astrocytes after severe stab injury showed some characteristics of A1 astrocytes, including massive increase in the expression level of *Gbp2* and *Serping1.* Surprisingly, after only 3 days of NeuroD1 infection, even before neuronal conversion, the reactive astrocytes lost most of the *Gbp2* and *Serping1* expression, together with a decrease of inflammatory marker Lcn2. Therefore, NeuroD1 expression inhibits the reactive response of astrocytes in the injury areas, as evident from the observation of less hypertrophic morphology and reduced GFAP and CSPG in NeuroD1-infected astrocytes.

It may be worth mentioning that a previous report found that mild stab injury in the brain resulted in less astrocytic proliferation (Bardehle et al., 2013). In the current study, we used a large needle (outer diameter 0.95 mm) to induce severe stab injury, resulting in clear tissue loss in the mouse motor cortex. Accordingly, we found a significant number of proliferating astrocytes in the injured areas. Therefore, different levels of neural injury or different disease models may result in different levels of astrocytic proliferation.

### Attenuation of neuroinflammation and restoration of BBB

It is unexpected when we detect a drastic decrease of microglia and neuroinflammatory factors in the injury areas following NeuroD1-mediated AtN conversion. If astrocytic and neuronal changes are directly associated with AtN conversion, the changes in microglia must be an indirect effect, an indication of non-cell autonomous impact on the microenvironment where the AtN conversion occurs. Microglia act as neuroprotective cells and play a pivotal role in immune defense in the brain by constantly surveying brain tissue to identify any potential damage for immediate repair. On the other hand, excessive activation of microglia often leads to secondary neuronal damage due to their secretion of neurotoxic cytokines, inflammatory factors, and reactive oxygen spices (Hu et al., 2015; Neher et al., 2013). Therefore, like reactive astrocytes, reactive microglia also exert both positive and negative effects on injured nerve tissue. How to maintain the immune response of microglia at a proper level is pivotal in neural repair. One major difficulty in controlling microglial response is to find out what is the best time window to stop their reactive responses once the immune defense system has already been turned on. In the current study, we were surprised to observe diminished microglial reactivity and a drastic reduction of inflammatory factors such as TNFa and interleukin 1β (IL-1β). Intriguingly, microglia that are in close contact with NeuroD1-infected astrocytes lost reactive properties, becoming iNOS negative with more ramified processes. This observation is in line with previous studies showing astrocytic modulation of microglial activity through a variety of factors including ATP, Ca^2+^, complement (C3), chemokines and plasma protein ORM2 (Clarner et al., 2015; Davalos et al., 2005; Jo et al., 2017; Lian et al., 2016; Schipke et al., 2002; Zamanian et al., 2012). Therefore, by converting reactive astrocytes into neurons, NeuroD1 treatment indirectly regulates the reactivity of microglia and reduces neuroinflammation.

Another indirect impact of AtN conversion is the restoration of the BBB integrity in NeuroD1-infected areas. Astrocytes actively participate in the maintenance of BBB integrity by physically contacting blood vessels with endfeet and secreting growth factors such as vascular endothelial growth factor A (VEGFA), basic fibroblast growth factor 2 (FGF2), interleukin 6 (IL-6), and glia-derived neurotrophic factor (GDNF) to promote angiogenesis (Obermeier et al., 2013). After stab injury, we found that the endfeet of reactive astrocytes were dissociated from the blood vessels and the water channel AQP4 was not concentrating at the endfeet anymore, suggesting a disruption of the interaction between astrocytes and blood vessels. This is consistent with previous finding that brain injury is accompanied with BBB breakdown (Shlosberg et al., 2010). Interestingly, after NeuroD1-mediated cell conversion, not only astrocytes became less reactive, but also astrocytic endfeet became re-associated with blood vessels and the AQP4 signal was relocated to the endfeet, suggesting a restoration of BBB. Such astrocytic endfeet change is indeed unexpected. The mislocalization of AQP4 signal in reactive astrocytes after brain injury is one of many indications that the reactive astrocytes cannot maintain normal structure and function in an injured environment (Fukuda and Badaut, 2012). Conversely, the reversal of AQP4 signal back to the endfeet is an indication that the neural injury has been alleviated and the astrocytes are returning to normal structure and function.

In summary, we demonstrate here that NeuroD1-mediated astrocyte-to-neuron conversion has broad impact on the microenvironment of injured neural tissue: it not only generates new neurons, reduces toxic A1 astrocytes, but also attenuates toxic M1 microglia and repairs blood vessels and BBB integrity. Importantly, the neural repairing effect of NeuroD1 starts very early, as early as 3-4 days post NeuroD1 infection and even before neuronal conversion. Our studies provide the proof-of-concept that glial scar can be reversed back to neural tissue through *in vivo* astrocyte-to-neuron conversion.

### Material and Methods

See supplementary material

## Acknowledgement

This work was supported by grants from National Institutes of Health (AG045656, MH083911), Alzheimer’s Association (ZEN-15-321,972), and Charles H. Smith Endowment Fund for Brain Repair from the Pennsylvania State University to G.C. We would like to thank all Chen Lab members for rigorous discussion throughout the past several years during the progress of this project.

## References

Allen, N.J., and Barres, B.A. (2009). Neuroscience: Glia - more than just brain glue. Nature 457, 675–677.

Anderson, M.A., Burda, J.E., Ren, Y., Ao, Y., O’Shea, T.M., Kawaguchi, R., Coppola, G., Khakh, B.S., Deming, T.J., and Sofroniew, M.V. (2016). Astrocyte scar formation aids central nervous system axon regeneration. Nature 532, 195–200.

Bardehle, S., Kruger, M., Buggenthin, F., Schwausch, J., Ninkovic, J., Clevers, H., Snippert, H.J., Theis, F.J., Meyer-Luehmann, M., Bechmann, I., et al. (2013). Live imaging of astrocyte responses to acute injury reveals selective juxtavascular proliferation. Nat Neurosci 16, 580–586.

Bradbury, E.J., Moon, L.D., Popat, R.J., King, V.R., Bennett, G.S., Patel, P.N., Fawcett, J.W., and McMahon, S.B. (2002). Chondroitinase ABC promotes functional recovery after spinal cord injury. Nature 416, 636–640.

Brambilla, R., Bracchi-Ricard, V., Hu, W.H., Frydel, B., Bramwell, A., Karmally, S., Green, E.J., and Bethea, J.R. (2005). Inhibition of astroglial nuclear factor kappaB reduces inflammation and improves functional recovery after spinal cord injury. J Exp Med 202, 145–156.

Bush, T.G., Puvanachandra, N., Horner, C.H., Polito, A., Ostenfeld, T., Svendsen, C.N., Mucke, L., Johnson, M.H., and Sofroniew, M.V. (1999). Leukocyte infiltration, neuronal degeneration, and neurite outgrowth after ablation of scar-forming, reactive astrocytes in adult transgenic mice. Neuron 23, 297–308.

Chung, I.Y., and Benveniste, E.N. (1990). Tumor necrosis factor-alpha production by astrocytes. Induction by lipopolysaccharide, IFN-gamma, and IL-1 beta. J Immunol 144, 2999–3007.

Clarner, T., Janssen, K., Nellessen, L., Stangel, M., Skripuletz, T., Krauspe, B., Hess, F.M., Denecke, B., Beutner, C., Linnartz-Gerlach, B., et al. (2015). CXCL10 triggers early microglial activation in the cuprizone model. J Immunol 194, 3400–3413.

Cregg, J.M., DePaul, M.A., Filous, A.R., Lang, B.T., Tran, A., and Silver, J. (2014). Functional regeneration beyond the glial scar. Exp Neurol 253, 197–207.

Davalos, D., Grutzendler, J., Yang, G., Kim, J.V., Zuo, Y., Jung, S., Littman, D.R., Dustin, M.L., and Gan, W.B. (2005). ATP mediates rapid microglial response to local brain injury in vivo. Nat Neurosci 8, 752–758.

Ferreira, A.C., Da Mesquita, S., Sousa, J.C., Correia-Neves, M., Sousa, N., Palha, J.A., and Marques, F. (2015). From the periphery to the brain: Lipocalin-2, a friend or foe? Prog Neurobiol 131, 120–136.

Fu, Y., Liu, Q., Anrather, J., and Shi, F.D. (2015). Immune interventions in stroke. Nat Rev Neurol 11, 524–535.

Fukuda, A.M., and Badaut, J. (2012). Aquaporin 4: a player in cerebral edema and neuroinflammation. J Neuroinflammation 9, 279.

Gascon, S., Murenu, E., Masserdotti, G., Ortega, F., Russo, G.L., Petrik, D., Deshpande, A., Heinrich, C., Karow, M., Robertson, S.P., et al. (2016). Identification and Successful Negotiation of a Metabolic Checkpoint in Direct Neuronal Reprogramming. Cell Stem Cell 18, 396–409.

Grande, A., Sumiyoshi, K., Lopez-Juarez, A., Howard, J., Sakthivel, B., Aronow, B., Campbell, K., and Nakafuku, M. (2013). Environmental impact on direct neuronal reprogramming in vivo in the adult brain. Nat Commun 4, 2373.

Guo, Z., Zhang, L., Wu, Z., Chen, Y., Wang, F., and Chen, G. (2014). In vivo direct reprogramming of reactive glial cells into functional neurons after brain injury and in an Alzheimer’s disease model. Cell Stem Cell 14, 188–202.

He, Z., and Jin, Y. (2016). Intrinsic Control of Axon Regeneration. Neuron 90, 437–451.

Heinrich, C., Bergami, M., Gascon, S., Lepier, A., Vigano, F., Dimou, L., Sutor, B., Berninger, B., and Gotz, M. (2014). Sox2-mediated conversion of NG2 glia into induced neurons in the injured adult cerebral cortex. Stem Cell Reports 3, 1000–1014.

Horn, K.P., Busch, S.A., Hawthorne, A.L., van Rooijen, N., and Silver, J. (2008). Another barrier to regeneration in the CNS: activated macrophages induce extensive retraction of dystrophic axons through direct physical interactions. J Neurosci 28, 9330–9341.

Hu, X., Leak, R.K., Shi, Y., Suenaga, J., Gao, Y., Zheng, P., and Chen, J. (2015). Microglial and macrophage polarization-new prospects for brain repair. Nat Rev Neurol 11, 56–64.

Jo, M., Kim, J.H., Song, G.J., Seo, M., Hwang, E.M., and Suk, K. (2017). Astrocytic Orosomucoid-2 Modulates Microglial Activation and Neuroinflammation. J Neurosci 37, 2878–2894.

Khakh, B.S., and Sofroniew, M.V. (2015). Diversity of astrocyte functions and phenotypes in neural circuits. Nat Neurosci 18, 942–952.

Koprivica, V., Cho, K.S., Park, J.B., Yiu, G., Atwal, J., Gore, B., Kim, J.A., Lin, E., Tessier-Lavigne, M., Chen, D.F., et al. (2005). EGFR activation mediates inhibition of axon regeneration by myelin and chondroitin sulfate proteoglycans. Science 310, 106–110.

Lian, H., Litvinchuk, A., Chiang, A.C., Aithmitti, N., Jankowsky, J.L., and Zheng, H. (2016). Astrocyte-Microglia Cross Talk through Complement Activation Modulates Amyloid Pathology in Mouse Models of Alzheimer’s Disease. J Neurosci 36, 577–589.

Liddelow, S.A., Guttenplan, K.A., Clarke, L.E., Bennett, F.C., Bohlen, C.J., Schirmer, L., Bennett, M.L., Munch, A.E., Chung, W.S., Peterson, T.C., et al. (2017). Neurotoxic reactive astrocytes are induced by activated microglia. Nature 541, 481–487.

Liu, Y., Miao, Q., Yuan, J., Han, S., Zhang, P., Li, S., Rao, Z., Zhao, W., Ye, Q., Geng, J., et al. (2015). Ascl1 Converts Dorsal Midbrain Astrocytes into Functional Neurons In Vivo. J Neurosci 35, 9336–9355.

Lu, P., Wang, Y., Graham, L., McHale, K., Gao, M., Wu, D., Brock, J., Blesch, A., Rosenzweig, E.S., Havton, L.A., et al. (2012). Long-distance growth and connectivity of neural stem cells after severe spinal cord injury. Cell 150, 1264–1273.

Nedergaard, M., Ransom, B., and Goldman, S.A. (2003). New roles for astrocytes: redefining the functional architecture of the brain. Trends Neurosci 26, 523–530.

Neher, J.J., Emmrich, J.V., Fricker, M., Mander, P.K., Thery, C., and Brown, G.C. (2013). Phagocytosis executes delayed neuronal death after focal brain ischemia. Proc Natl Acad Sci U S A 110, E4098–4107.

Niu, W., Zang, T., Zou, Y., Fang, S., Smith, D.K., Bachoo, R., and Zhang, C.L. (2013). In vivo reprogramming of astrocytes to neuroblasts in the adult brain. Nat Cell Biol 15, 1164–1175.

Norden, D.M., Fenn, A.M., Dugan, A., and Godbout, J.P. (2014). TGFbeta produced by IL-10 redirected astrocytes attenuates microglial activation. Glia 62, 881–895.

Obermeier, B., Daneman, R., and Ransohoff, R.M. (2013). Development, maintenance and disruption of the blood-brain barrier. Nat Med 19, 1584–1596.

Ojala, D.S., Amara, D.P., and Schaffer, D.V. (2015). Adeno-associated virus vectors and neurological gene therapy. Neuroscientist 21, 84–98.

Peron, S., Droguerre, M., Debarbieux, F., Ballout, N., Benoit-Marand, M., Francheteau, M., Brot, S., Rougon, G., Jaber, M., and Gaillard, A. (2017). A Delay between Motor Cortex Lesions and Neuronal Transplantation Enhances Graft Integration and Improves Repair and Recovery. J Neurosci 37, 1820–1834.

Pineau, I., Sun, L., Bastien, D., and Lacroix, S. (2010). Astrocytes initiate inflammation in the injured mouse spinal cord by promoting the entry of neutrophils and inflammatory monocytes in an IL-1 receptor/MyD88-dependent fashion. Brain Behav Immun 24, 540–553.

Rivetti di Val Cervo, P., Romanov, R.A., Spigolon, G., Masini, D., Martin-Montanez, E., Toledo, E.M., La Manno, G., Feyder, M., Pifl, C., Ng, Y.H., et al. (2017). Induction of functional dopamine neurons from human astrocytes in vitro and mouse astrocytes in a Parkinson’s disease model. Nat Biotechnol 35, 444–452.

Schipke, C.G., Boucsein, C., Ohlemeyer, C., Kirchhoff, F., and Kettenmann, H. (2002). Astrocyte Ca2+ waves trigger responses in microglial cells in brain slices. FASEB J 16, 255–257.

Sekiya, T., Holley, M.C., Hashido, K., Ono, K., Shimomura, K., Horie, R.T., Hamaguchi, K., Yoshida, A., Sakamoto, T., and Ito, J. (2015). Cells transplanted onto the surface of the glial scar reveal hidden potential for functional neural regeneration. Proc Natl Acad Sci U S A 112, E3431–3440.

Shlosberg, D., Benifla, M., Kaufer, D., and Friedman, A. (2010). Blood-brain barrier breakdown as a therapeutic target in traumatic brain injury. Nat Rev Neurol 6, 393–403.

Silver, J. (2016). The glial scar is more than just astrocytes. Exp Neurol 286, 147–149.

Silver, J., and Miller, J.H. (2004). Regeneration beyond the glial scar. Nat Rev Neurosci 5, 146–156.

Sivasankaran, R., Pei, J., Wang, K.C., Zhang, Y.P., Shields, C.B., Xu, X.M., and He, Z. (2004). PKC mediates inhibitory effects of myelin and chondroitin sulfate proteoglycans on axonal regeneration. Nat Neurosci 7, 261–268.

Sofroniew, M.V. (2015). Astrocyte barriers to neurotoxic inflammation. Nat Rev Neurosci 16, 249–263.

Su, Z., Niu, W., Liu, M.L., Zou, Y., and Zhang, C.L. (2014). In vivo conversion of astrocytes to neurons in the injured adult spinal cord. Nat Commun 5, 3338.

Toft, A., Scott, D.T., Barnett, S.C., and Riddell, J.S. (2007). Electrophysiological evidence that olfactory cell transplants improve function after spinal cord injury. Brain 130, 970–984.

Torper, O., Ottosson, D.R., Pereira, M., Lau, S., Cardoso, T., Grealish, S., and Parmar, M. (2015). In Vivo Reprogramming of Striatal NG2 Glia into Functional Neurons that Integrate into Local Host Circuitry. Cell Rep 12, 474–481.

Torper, O., Pfisterer, U., Wolf, D.A., Pereira, M., Lau, S., Jakobsson, J., Bjorklund, A., Grealish, S., and Parmar, M. (2013). Generation of induced neurons via direct conversion in vivo. Proc Natl Acad Sci U S A 110, 7038–7043.

Volterra, A., and Meldolesi, J. (2005). Astrocytes, from brain glue to communication elements: the revolution continues. Nat Rev Neurosci 6, 626–640.

Wanner, I.B., Anderson, M.A., Song, B., Levine, J., Fernandez, A., Gray-Thompson, Z., Ao, Y., and Sofroniew, M.V. (2013). Glial scar borders are formed by newly proliferated, elongated astrocytes that interact to corral inflammatory and fibrotic cells via STAT3-dependent mechanisms after spinal cord injury. J Neurosci 33, 12870–12886.

Witcher, K.G., Eiferman, D.S., and Godbout, J.P. (2015). Priming the inflammatory pump of the CNS after traumatic brain injury. Trends Neurosci 38, 609–620.

Yiu, G., and He, Z. (2006). Glial inhibition of CNS axon regeneration. Nat Rev Neurosci 7, 617–627.

Zamanian, J.L., Xu, L., Foo, L.C., Nouri, N., Zhou, L., Giffard, R.G., and Barres, B.A. (2012). Genomic analysis of reactive astrogliosis. J Neurosci 32, 6391–6410.

